# Brain Templates for Chinese Babies from Newborn to Three Months of Age

**DOI:** 10.1101/2023.06.05.543553

**Authors:** Xiujuan Geng, Peggy HY Chen, Hugh S Lam, Winnie CW Chu, Patrick CM Wong

## Abstract

The infant brain develops rapidly and this area of research has great clinical implications. Neurodevelopmental disorders such as autism and developmental delay have their origins, potentially, in abnormal early brain maturation. Searching for potential early neural markers requires *a priori* knowledge about infant brain development and anatomy. One of the most common methods of characterizing brain features requires normalization of individual images into a standard stereotactic space and conduct of group-based analyses in this space. A population representative brain template is critical for these population-based studies. Little research is available on constructing brain templates for typical developing Chinese infants. In the present work, a total of 112 babies from 6 to 98 days of age were included with high resolution structural magnetic resonance imaging scans. T1-weighted and T2-weighted templates were constructed using an unbiased registration approach for babies from newborn to 3 months of age. Age-specific templates were also estimated for babies aged at 0, 1, 2 and 3 months old. Then we conducted a series of evaluations and statistical analyses over whole tissue segmentations and brain parcellations. Compared to the use of population mismatched templates, using our established templates resulted in lower deformation energy to transform individual images into the template space and produced a smaller registration error, i.e., smaller standard deviation of the registered images. Significant volumetric growth was observed across total brain tissues and most of the brain regions within the first three months of age. The total brain tissues exhibited larger volumes in baby boys compared to baby girls. To the best of our knowledge, this is the first study focusing on the construction of Chinese infant brain templates. These templates can be used for investigating birth related conditions such as preterm birth, detecting neural biomarkers for neurological and neurodevelopmental disorders in Chinese populations, and exploring genetic and cultural effects on the brain.

## Introduction

The infant brain is constantly changing in size and morphology due to its rapid development. Standard maps of the infant brain, containing *a priori* knowledge about neural anatomy would help neuroscientists and clinicians in their investigation of neurodevelopmental issues. A large portion of neurodevelopmental disorders, such as schizophrenia, autism, and ADHD have their potential origins in abnormal early brain maturation (Bale et al., 2010; Visser et al., 2016; Owen & O’Donovan, 2017). Emerging evidence also suggests that the early brain imaging markers are associated with later behavioral and cognitive outcomes (Woodward et al., 2006; Gabrieli et al., 2015; Ullman et al., 2015; Short et al., 2019). Investigation of early brain development provides a unique approach for the discovery of neural basis of mental disorders and behavioral disabilities. Infant brain templates and atlases serve as important references for the studies of early brain development with magnetic resonance imaging (MRI). Normalization-based image analysis is one of the most common methods for quantification of the neuroimaging data among populations. Individual images are spatially normalized into the standard stereotactic space, and the following analyses are conducted in this space. Therefore, appropriate infant brain templates are critical for estimating neural measures in babies.

The existing infant brain templates and atlases, such as UNC Infant atlases (Shi et al., 2011), JHU neonate atlas (Oshi et al., 2011) and the spatio-temporal neonatal atlas (Makropoulos et al., 2016), are mostly based on images obtained from Caucasian babies, which may not be suitable for studying brain development of Chinese infants due to neuroanatomical differences related to genetic and environmental factors. Although there is a lack of investigations comparing infant brains between Western and Chinese populations, morphological differences have been reported between Caucasian and East Asian adults when measuring brain volumetric and cortical features such as regional volumes, surface area and cortical thickness in temporal, frontal and parietal brain regions (Chee et al., 2011; Tang et al., 2018; Huang et al., 2019). Moreover, the overall brain morphological features, e.g., length, width, and total brain and tissue volumes, differ between US and Chinese children and adolescents (Xie et al., 2015). A study on the construction of Chinese children also revealed anatomical differences in frontal and parietal regions between the two populations (Zhao et al 2019). Furthermore, when considering developmental disorders, such as dyslexia, unique disruptions in the brains of Chinese children have been evidenced compared to Western children (Siok et al., 2004). Based on the findings of distinct brain morphology in children and adults from different ethnicities with various genetic and cultural factors, we postulate that there are also structural differences between the Western and Chinese infant brains.

These neural variations across cultures may lead to a mismatch when the template constructed from one population is used on the images from another, and the template is considered as a population mismatched template. A study on the brain templates of adults from Western and East Asian populations has reported different variabilities in the deformation degree, which measures how much spatial normalization an individual needs to transform to the template image. And the brain regions exhibiting distinct transforming patterns are mostly located in language areas, i.e., supramarginal gyri and inferior frontal gyri (Yang et al., 2020). Furthermore, image processing performance, such as accuracy in brain segmentation and registration, will be reduced when using the population mismatched templates. Due to the lack of infant brain templates based on Chinese infants, there is a need to establish an unbiased and population-specific infant brain template, which may be more suitable to characterize the early neural development of Chinese babies.

In this work, we focused on the construction of infant brain templates using T1-weighted and T2-weighted MRI data from over one hundred Chinese babies aged between newborn and three months old. By comparing to the neonatal template established with Western babies (Shi et al., 2011), we evaluated and demonstrated the advantage of the use of our population-matched templates. With the population matched templates, we also examined the overall and regional brain developmental trajectories within the population.

## Methods

### Subjects

Participants were recruited from the Prince of Wales Hospital (PWH) at Hong Kong and from the community through media interviews and use of social networks (Facebook and instant messenger groups). Permission for MRI was granted by the joint Chinese University of Hong Kong – New Territories East Cluster Clinical Research Ethics Committee (ethics number 2020.144). Written informed consent was obtained from parents or legal guardians of all participants according to the principles explained in the Declaration of Helsinki, and the rights of these participants were protected accordingly. Babies included in this study were born between 32-40 weeks gestation and had passed the hearing screening. The exclusion criteria included: birth asphyxia, major birth injuries, hypoxic-ischemic injury, significant growth restriction (<5^th^ percentile) and other medical conditions that affect language and neurocognitive developments.

All MRI scans needed to pass quality control to be included in this study. The detailed quality control procedure is described below. First, the anatomical images of all subjects were carefully screened by an experienced pediatric radiologist (WCWC) to exclude any incidental abnormalities, including developmental anomaly or intracranial space-occupying mass lesions such as neuroepithelial cysts or arachnoid cysts. Then, careful visual inspections of the scans were conducted using a protocol as follows: full coverage of the brain; no obvious motion artifacts; and reasonable signal to noise ratio. A total number of 98 T2-weighted, and 86 T1-weighted images from 112 babies were included for template building (see Table 1). Babies ranged between 6 and 98 days of corrected postnatal age with a mean age of 47.04 days. All subjects were healthy, typically developing, moderate to late preterm or early term babies, with no obvious congenital, neurological or physical abnormalities or impairments. Only one infant required invasive mechanical ventilation for two hours. The subjects included in our studies were born at gestational age from 32 weeks to 39 weeks +4 days. None of them was admitted to a Neonatal Intensive Care Unit (NICU).

**Table 1.** Participant characteristics.

|  | Range | Mean | Number (%) |
| --- | --- | --- | --- |
| **Gestational age at birth (days)** | 32w to 39w + 4 d | 36.3 d |  |
| **Postmenstrual age (days)** | 16-44 days | 29.93 d |  |
| **Postnatal age (days)** | 21 to 133 days | 72.54 d |  |
| **Corrected postnatal age (days)** | 6 to 89 days | 47.04 d |  |
| **Gestational age (weeks)** | 32w to 39w + 4 d | 36w + 2d |  |
| **Female, n** |  |  | 52 (46.25) |
| **Birthweight (g)** | 1280-4780 | 2586 |  |
| **Resuscitation &/or requirement of oxygen and ventilation support at birth** |  |  | 19 (17%) |

### Image Acquisition

Subjects were scanned during natural sleep without sedation on a Siemens Magnetom Prisma 3T scanner in PWH at Hong Kong. Before the scans, infants were wrapped up and soft pads and/or earmuffs were used to securely cover and protect the ears from scanner noise. Ear plugs were also inserted in the ears of babies who were sensitive to noise. A parent or guardian of the baby sat inside the scanner room to observe whether the baby was sleeping throughout the scan. Higher-resolution T1-weighted (T1w) and T2-weighted (T2w) images were acquired by a 16-channel pediatric coil with MPRAGE and 3D SPACE sequences in the sagittal plane respectively. Main imaging parameters of T1w were as follows: repetition time (TR)= 1800 ms; echo time (TE)=2.82ms; flip angle=8°; FOV= 180×180×128 mm; voxel size=0.8×0.8×0.8 mm; acquisition matrix=224×224×160. The T2w parameters were: TR=3200 ms; TE= 564 ms; flip angle=T2 var; FOV= 205×205×128 mm; voxel size= 0.8×0.8×0.8 mm; acquisition matrix=256×256×160. The acquisition time was 4:05 mins for T1w and 4:50 mins for T2w.

### Template Construction

#### Image Preprocessing

All MRI scans were first preprocessed using the steps described below. i) Images were corrected for intensity inhomogeneity using N3 intensity non-uniformity method (Sled et al., 1998). ii) The inputs were resampled with 1 mm^3^, reformats to UCHAR. iii) A skull stripping method HD-BET (Isensee et el., 2019) was conducted to extract data related to the brain for T2w images, and an infant MRI skullstripping method with convolutional neural networks (Jog et al., 2019) was applied for T1w images. iv) The image intensity of all images was normalized by a histogram-matching method (Nyúl et al., 2000) for T1w images and T2w images.) A hierarchical linear registration of each scan to the UNC Neonate (UNC_Neo) template was conducted as follows: the data from the original space was first rigidly aligned to the UNC_Neo template with 6 parameters (3 rotation and 3 translation), then the average of the rigidly aligned data was computed as the initial rigid template, and the data from the original space was affine aligned to the initial rigid template with 12 parameters. The affine aligned data was averaged to obtain the initial affine template.

#### Unbiased Template Construction

An unbiased template construction procedure was conducted to generate the infant brain atlas. As mentioned above, the preprocessed images were first averaged as the initial reference target template. Then the deformable symmetric image normalization (SyN) method in ANTs was conducted to warp each individual image into this initial template (Avants et al., 2009; Avants et al., 2011). This nonlinear registration procedure was repeated three times (based on empirical evidence) to obtain an optimal template. This template creation process was conducted for all images to generate an overall template for infants.

Due to the fast maturation speed right after birth (Gilmore et al., 2012; Holland et al., 2014), it is necessary to construct a set of templates, with each one corresponding to a specific and narrow age range. Therefore, we also constructed a series of age-specific templates for babies aged at 0, 1, 2 and 3 months, named T_0, T_1, T_2, T_3, respectively. When constructing these templates, subjects included for each template were the ones with corrected age rounded to that specific month (Table 2): T_0 for 0-month (6-13 days of age); T_1 for 1-month (16-44 days); T_2 for 2-month (45-74 days); and T_3 for 3-month old (75-89 days) (Table 2). The original images within one age group were averaged, then rigidly aligned to the initial average to obtain the rigid average, and affine aligned to the rigid-aligned average to get the initial affine template. Then the nonlinear registration was performed three times to obtain the age-specific template. The aforementioned template construction process was conducted separately on T2w and T1w images to obtain templates for each modality. Therefore, a total of ten templates were built, five templates (four for each of the four age groups and one for overall) based on T2w images and five based on T1w images.

**Table 2.**
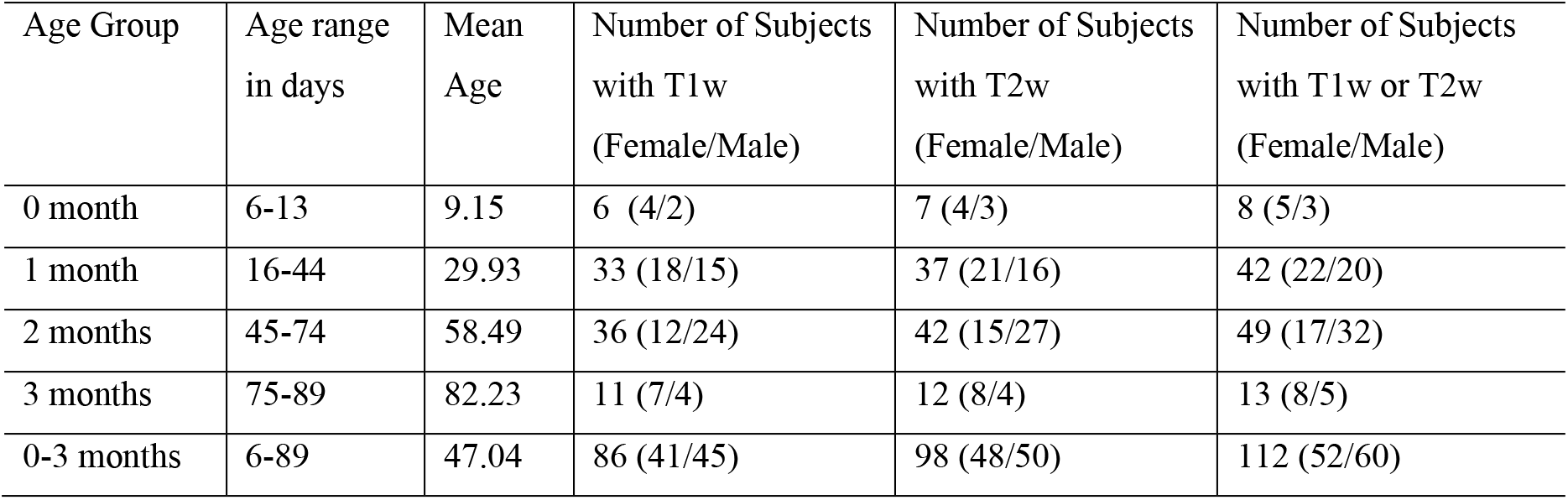
Demographic and scan availability information of babies included in each age group.

| Age Group | Age range<br>in days | Mean<br>Age | Number of Subjects<br>with T1w<br>(Female/Male) | Number of Subjects<br>with T2w<br>(Female/Male) | Number of Subjects<br>with T1w or T2w<br>(Female/Male) |
| --- | --- | --- | --- | --- | --- |
| 0 month | 6-13 | 9.15 | 6 (4/2) | 7 (4/3) | 8 (5/3) |
| 1 month | 16-44 | 29.93 | 33 (18/15) | 37 (21/16) | 42 (22/20) |
| 2 months | 45-74 | 58.49 | 36 (12/24) | 42 (15/27) | 49 (17/32) |
| 3 months | 75-89 | 82.23 | 11 (7/4) | 12 (8/4) | 13 (8/5) |
| 0-3 months | 6-89 | 47.04 | 86 (41/45) | 98 (48/50) | 112 (52/60) |

Besides the templates, we have also propagated the tissue segmentation (segmentations for gray matter (GM), white matter (WM), and cerebrospinal fluid (CSF)) and 90 regions of parcellation based on automated anatomical labelling (AAL) (Tzourio-Mazoyer et al., 2002) in the UNC_Neo space to our population-specific unbiased space by registering the UNC_Neo template to our templates. Notably, our template construction procedure can be used to generate customized infant and pediatric templates according to different age ranges of interests.

### Template Evaluation

To estimate whether the constructed templates are superior, we chose the UNC neonate template for comparison. We randomly chose 90% of the images for template building, and the remaining 10% for testing, and repeated this procedure ten times. The testing images were registered to the validation template (constructed by 90% of the images). Then the standard deviation of the registered images on each voxel was calculated and compared using the UNC_Neo and the validation template with the same registration methods. The transformation was also compared by computing the determinant of the Jacobian of the transformation fields. To quantify the metrics according to different brain areas, we propagated the AAL parcellation, which was in the UNC_Neo template space, to the validation template space by registering the UNC_Neo to it with SyN. Then the average of the determinant of the Jacobian maps JH(x) (Eq. (1)) and standard deviation maps STD(x) (Eq. (2)) over each ROI was computed and compared between using different templates. A smaller degree of displacements from warping an image to the template suggests a more population-representative template. Smaller standard deviation indicates smaller registration error using the corresponding template.

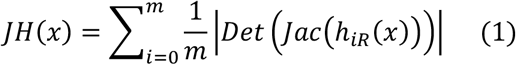

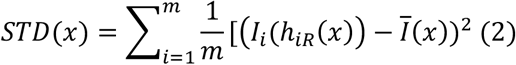

### Analysis of Age-Related Development Using the Unbiased Template

To illustrate the usage of the unbiased template, we have also analyzed the regional volumetric development within the Chinese babies aged between 0 to 3 months. A general linear model was used to examine the relationship between brain volume and the age at scan controlling the gestational age (GA) and sex: volume = GA + Sex + Age. This model was used to examine the development of overall tissue volume and the regional volume as a function of age in days. All analyses were conducted in R.

## Results

### Structural Templates, Tissue Segmentation, and Brain Parcellation

The T2-weighted and T1-weighted templates were constructed based on a total of 112 babies aged 6-98 days, 98 for T2-weighted and 86 for T1-weighted. Figure 1(a) shows the three (axial, sagittal and coronal) views of the overall brain template based on T1w and T2w images, and templates of each age group. The overall size growth could be observed from newborn to 3 months of age. The brain tissue segmentation and parcellation in the standard template space, for all babies, and babies of 0, 1, 2 and 3 months of age, were displaced in Figure 1(b), which were obtained by warping the labels in the UNC_Neo space to our constructed template space.

**Figure 1.**
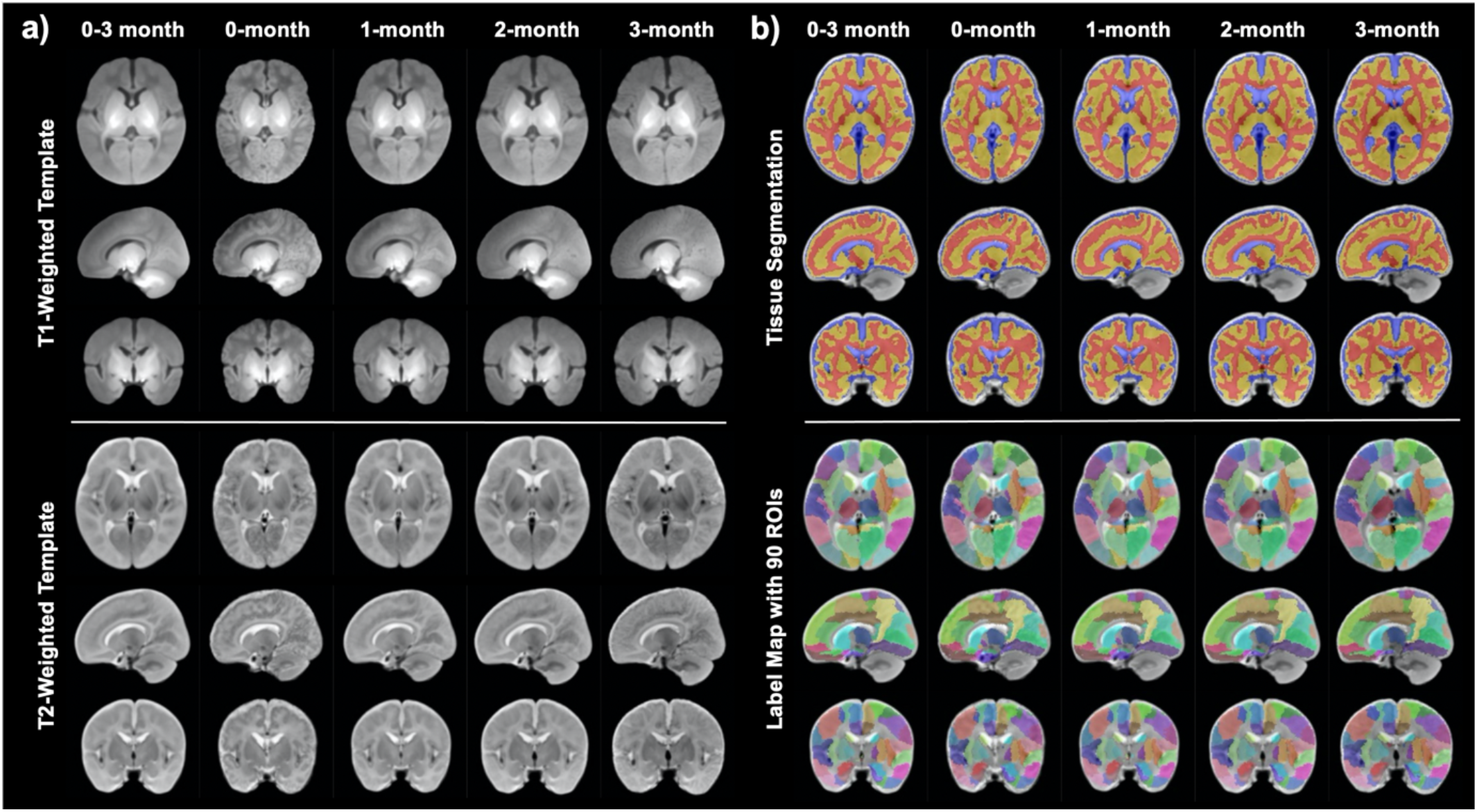
The unbiased population specific templates and the corresponding tissue segmentation and brain parcellation. (a) The templates using T1w (top panel) and T2w (bottom panel) images for all babies aged from 0 to 3 months, and for babies of 0, 1, 2, and 3 months of age. (b) Tissue segmentation (red: white matter; yellow: gray matter; blue: CSF), and label maps with 90 ROIs (each color represents a ROI) in the corresponding template space.

### Evaluation of the Unbiased Population Specific Templates

When comparing our established population-specific templates with the existing newborn infant brain template UNC_Neo, the Jacobian and standard deviation maps were calculated to quantify the advantage of using the population representative template. Smaller Jacobian determinants when using the population-matched template were observed in various brain areas especially in the amygdala and parahippocampus, globus pallidus, and medial prefrontal regions (Figure 2(a) & (b)). Forty-seven out of 90 ROIs showed significant reduced Jacobian determinants (Bonferroni corrected) when using the unbiased and population matched template (Figure 3, Table S1). Moreover, the use of the population-matched template also exhibited smaller standard deviation values compared to the use of the population-mismatched template in several brain regions including the parahippocampus, amygdala and anterior cingulate (Figure 2(c) & (d)). Sixteen ROIs showed significantly smaller standard deviations (Bonferroni corrected) when using the unbiased template (Figure 4, Table S2). Findings suggest that the unbiased population-specific template may provide more representative standard space for spatially normalizing individual images in terms of smaller warping energy (smaller Jacobian determinants) and more accurate alignment into the standard space (smaller standard deviations).

**Figure 2.**
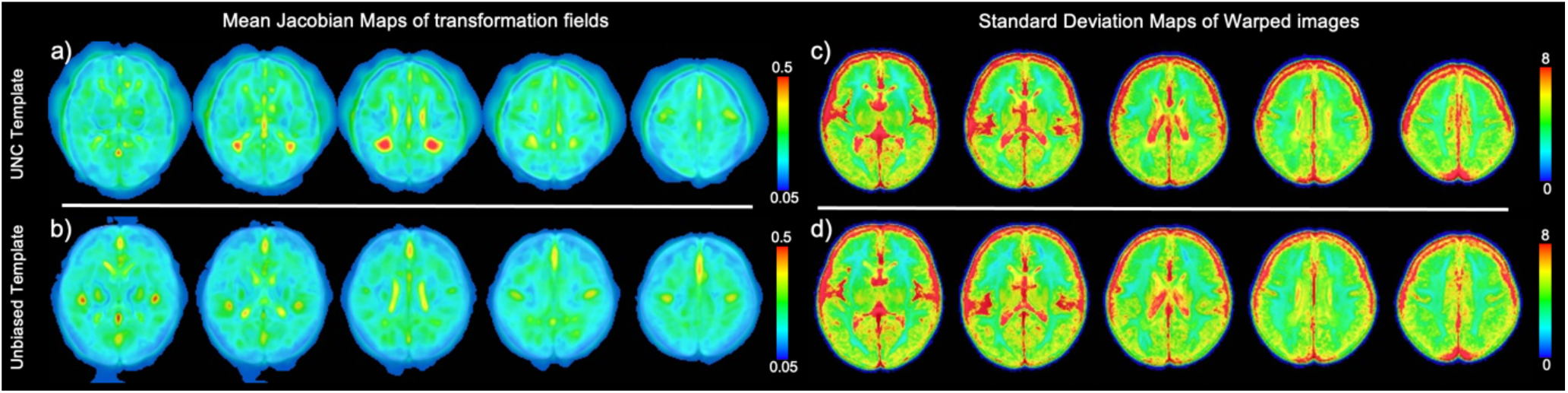
Comparison between the use of the population mismatched (UNC_Neo) and matched templates. (a) The average of the Jacobian determinant maps of the transformation fields. (b) The average of the standard deviation maps of the warped images to the template space.

**Figure 3.**
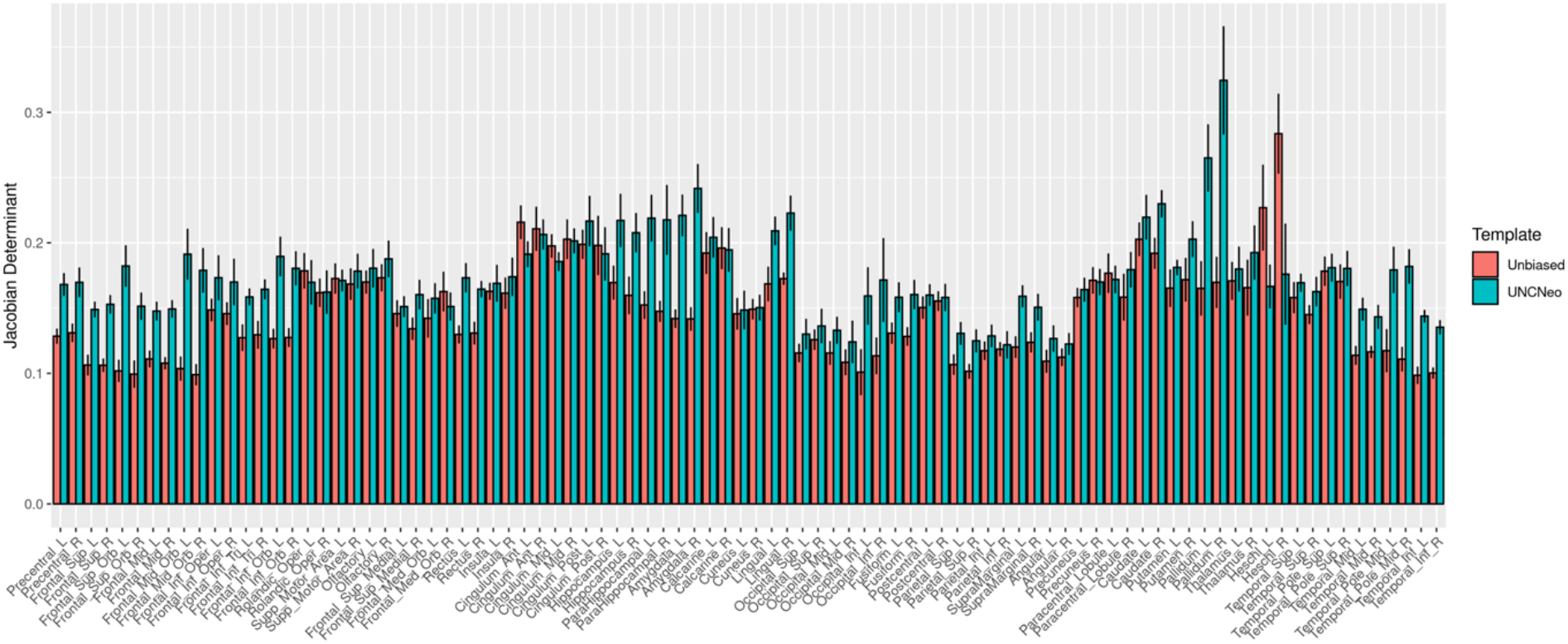
Plot of the average Jacobian determinant maps of the transformation fields across 90 ROIs when transforming individuals to the unbiased population specific template CUHK_Neo and transforming to population mismatched template UNC_Neo. The majority of the regions show smaller Jacobian determinants.

**Figure 4.**
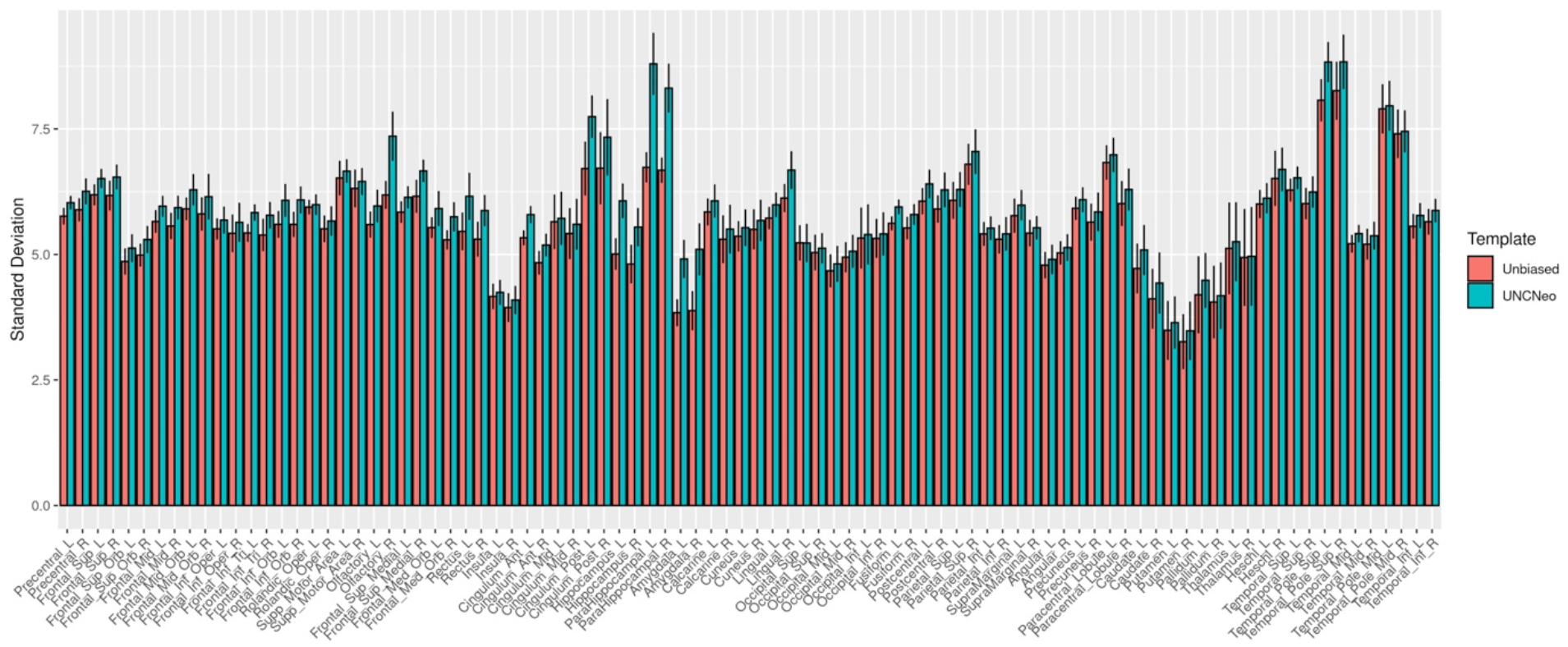
Plot of the standard deviation of the registered images to the unbiased population matched template CUHK_Neo and population mismatched template UNC_Neo across 90 brain regions. The majority of the regions show smaller standard deviations when registering to the CUHK_Neo template.

### Analyses of Age Related Development

We analyzed the developmental patterns of overall and regional volumetric growth within babies aged 0 to 3 months. Figure 5 plots the tissue volumes vs. age in days. All three tissue volumes were significantly positively correlated with age after correcting for GA with all corrected ps <0.001. And the development of the 90 regional volumes over age was plotted in Figure 6. Note that, except the volumes of four regions, bilateral posterior cingulate and caudate, all other volumes show significant growth over age (all ps < 0.05 after Bonferroni correction) (Table S3). In order to examine the overall brain volumes, we have also calculated the total brain volume and total tissue volumes for all babies and babies within one, two and three months of age (Table 3). The total cerebral volumes were 423.6±37.5 cm^3^, 486.2±47.3 cm^3^ and 524.1±39.3 cm^3^ for babies within one, two and three months of age, respectively.

**Table 3.**
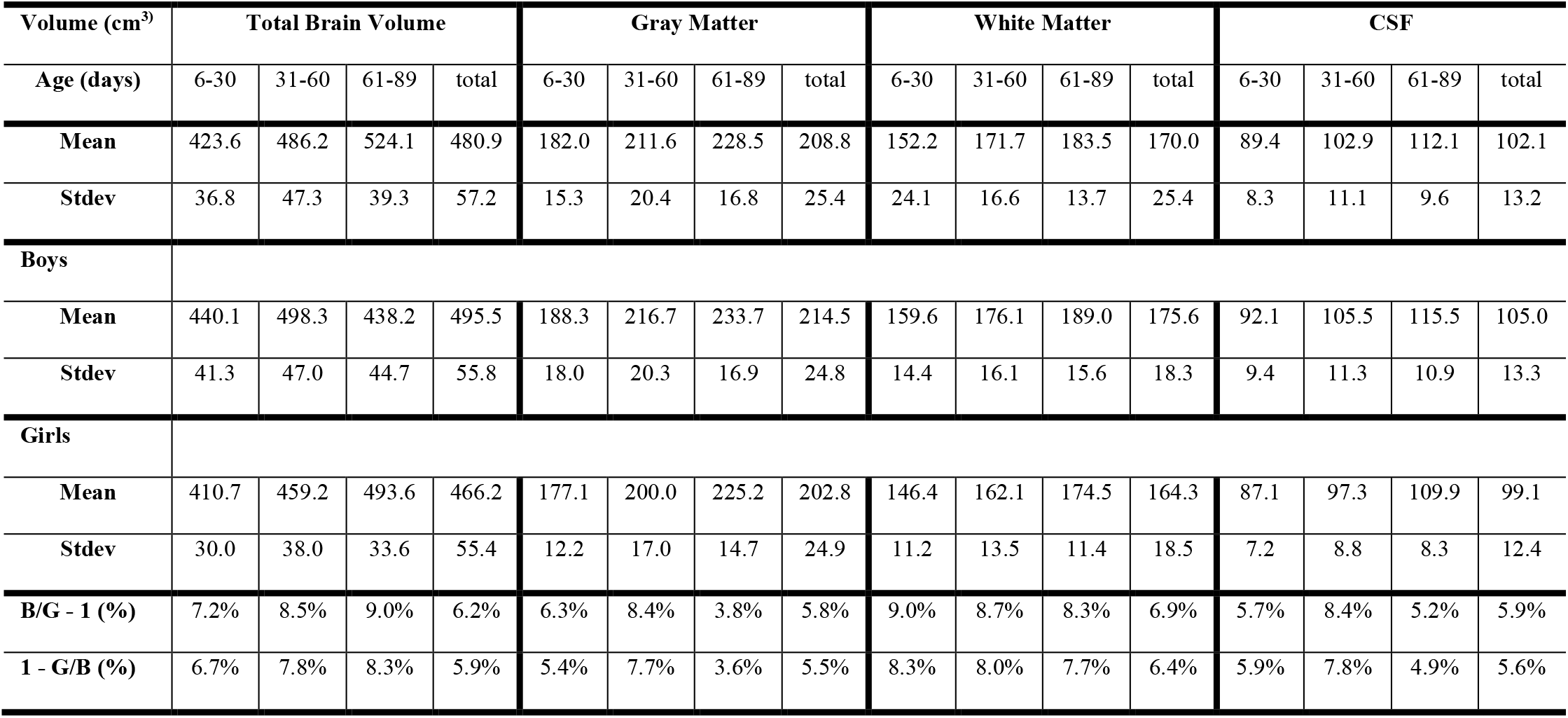
Brain volumetric measures at different postnatal ages.

**Figure 5.**
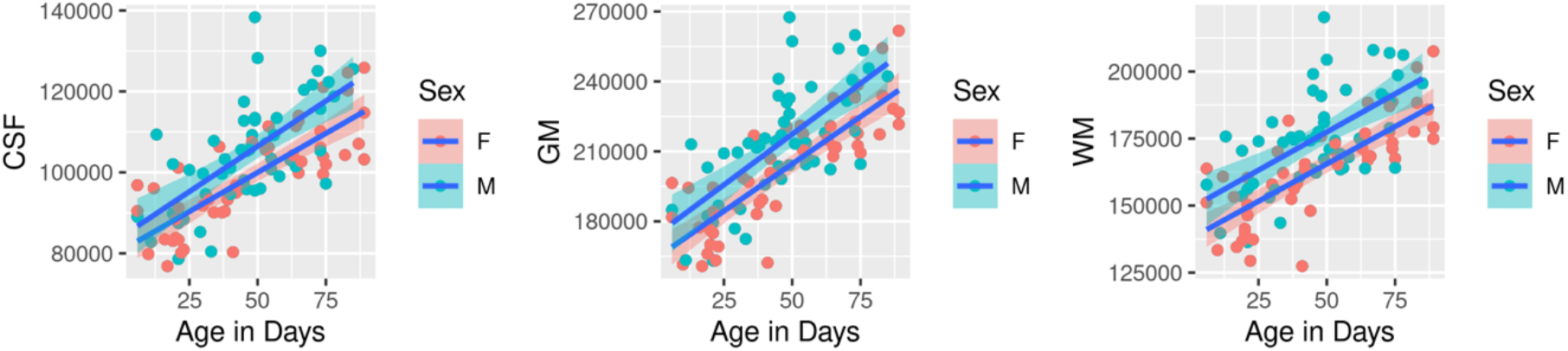
Volume development of brain tissues of cerebrospinal fluid (CSF), gray matter (GM) and white matter (WM) over age at the MRI scan.

**Figure 6.**
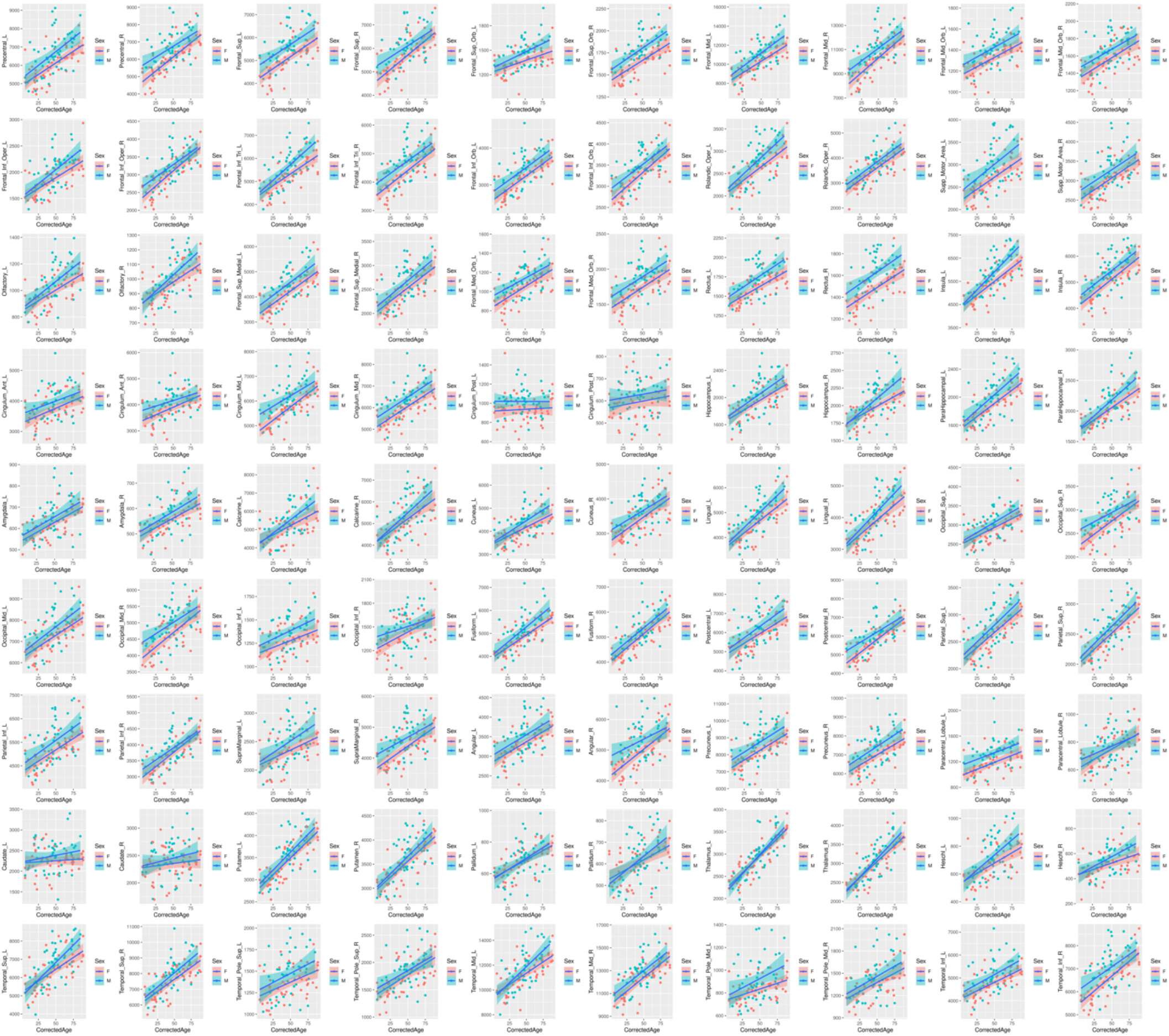
Regional volume development of 90 ROIs over age at the MRI scan. All volumes except for the volumes of bilateral posterior cingulate and caudate showed significant growth (corrected ps < 0.05).

As a post-hoc analysis, we also checked whether the overall tissue volumes and regional volumes were affected by sex or not. Significant sex effect on the three tissue volumes were found where the baby boys had greater volume than baby girls, with corrected p=0.0034 for CSF; p=0.00080 for GM; and p=0.00011 for WM. On average, boys showed 5.8%, 6.9%, and 5.9% larger volume of GM, WM, and CSF, respectively, compared to those of girls within 3 months of age (Table 3). When checking on regional volume, the following areas showed significant effect of sex (corrected ps < 0.05) (Table S3): bilateral rectus, right superior orbital frontal cortex, left superior frontal cortex, right precentral gyrus, left middle cingulum, left paracentral lobule, right medial orbital frontal cortex and left insula. Boys showed larger volumes in these frontal and prefrontal cortical regions. When controlling the total brain volume, no significant regional volume difference existed between boys and girls.

## Discussion

In the current study, we have constructed a set of Chinese infant templates using 112 babies aged 0-3 months with an average age of 47.04 days using T1w and T2w images. We also constructed age-specific templates for infants aged at 0, 1, 2 and 3 months old. The tissue segmentations and 90 regions of parcellations were estimated by propagating them from the UNC_Neo template space. Smaller warping energy was required to spatially normalize individuals to our established population-matched template compared to warping to the neonate template of Western infants. Transforming to our constructed templates also was associated with smaller registration error in terms of smaller standard variation of the warped images. We have also observed significant volumetric development for the overall brain tissues, and all regions except bilateral posterior cingulate cortex and caudate, within the first three months of age. The brain tissues of GM, WM and CSF, and several prefrontal and frontal regions also exhibit sex effects, where baby boys show larger volumes than baby girls.

To the best of our knowledge, this is the first attempt to construct Chinese infant templates. There is a great need of population-matched templates for the study of infant brain development and related neurodevelopmental disorders. Although very few studies directly compare brain morphology between Western and Chinese babies, evidence shows that there are differences in overall brain size and shape in children and adolescents (Xie et al., 2015) and regional volumes and cortical features in children (Zhao et al 2019) and adults (Tang et al., 2018; Huang et al., 2019). All these reported findings indicate that the baby brains would also differ in morphology globally and regionally which might be due to genetic and/or cultural and environmental factors. Moreover, the use of population mis-matched templates may cause larger errors in analysis (Yang et al., 2020). Our findings of larger deformation energy and bigger registration error when using population mis-matched templates provide direct evidence for the need of population-specific templates for Chinese babies.

It has been found that amongst Chinese children, incidences of neurodevelopmental disorders including ASD (Wang et al., 2018), ADHD (Liu et al., 2018), and language disorder (Wu et al., 2023) are substantial. The knowledge of early brain morphology in babies provides valuable insights to investigate the neural developmental deviations in populations under various conditions. The existing infant brain templates, such as the UNC Infant atlases (Shi et al., 2011; Oshi et al., 2011; Makropoulos et al., 2016) are mostly based on images obtained from Caucasian populations which may not be suitable for the investigation of the brains of Chinese infants due to neuroanatomical differences related to genetic and environmental factors. Therefore, it is time to construct templates and atlases based on Chinese infants to facilitate the investigation of brain mechanisms underlying neurodevelopmental disorders in this population.

There has been evidence to associate cerebral volumes with impaired neurocognitive outcomes (Lind et al., 2011; Van Kooij et al., 2012). However, there is lack of comparison of brain morphology from various cultures and ethnicities. It is worth investigating the neural volumetric and anatomical differences of babies across cultures and ethnicities, and whether the diversity comes from nature or nurture (Ge et al., 2023). A study on a cohort of Chinese babies from the NICU reported a total brain volume of 433±40.61 cm^3^ at the postmenstrual age (PMA) from 40 to 43 weeks of age, with a confidence interval of [422.53, 444.29] (Wang et al., 2018). The total brain volume in our study was 422 cm^3^ for babies within 1 month in corrected postnatal age, which is comparable to Wang et al.’s work. Another meta-analysis on the total brain volume of preterm babies, mostly Caucasian, reported a pooled value of 399 cm^3^ (standard deviation of 57) at the term equivalent age of 41weeks of postmenstrual age. Until now, however, we have not been able to make legitimate comparisons between Western and Eastern babies, mainly due to the lack of studies and available cohorts focusing on the investigation of babies in the populations. Our work of recruiting a Chinese baby cohort and establishing the brain template is a critical step toward this research direction.

Sex differences in total brain volume have been evidenced to be present at birth and continue to exist throughout postnatal development (See a review from Eliot et al., 2021). Our study also suggested that baby boys present with 6.2% larger brain volume compared with baby girls, which lies within the reported volume difference in Western countries by Knickmeyer and colleagues (suggesting that newborn boys have 5% larger brain size) (Knickmeyer et al., 2014) and Dean et al.’ (boys with 8% larger at 1 month old) (Dean et al., 2018). However, when controlling for the total brain volume, we did not find significant difference in regional volumes between boys and girls. Due to the unequal prevalence of psychiatric and neurological disorders, such as autism and depression, exhibited between males and females, researchers tend to assume that there are pre-existing sex differences in the brain. However, the sex “dimorphism” is still an inconclusive statement (Eliot et al., 2021). No sex differences on cortical parameters have been reported in babies from newborn to two years of age (Geng et al., 2017). A recent study reported greater thickness in male adults within both Chinese and American populations, but the regions showing differences do not overlap between the two populations (Yang et al., 2020b). All these suggest that more studies are needed to elucidate any possible sex differences in the brain, which may be population-specific, and influenced by a combination of genetic and environmental factors.

Since the baby brain experiences rapid growth in size and function due to the fast neural maturation during the early age of postnatal period (Geng et al., 2012; Lyall et al., 2014; Holland et al., 2014; Geng et al., 2017), it is becoming clear that a single brain template is not sufficient to characterize the time-dependent changes (Oishi et al., 2019; Chen et al., 2022). In the current work, we have split subjects into age groups of 0, 1, 2, and 3 months and established age-specific templates for each age group. With the growing number of collected babies, our future work will focus on extending the construction of spatiotemporal brain templates and atlases for Chinese babies.

For template construction, besides the use of either T1w or T2w data, it is preferred to combine both of them together with other imaging modalities such as diffusion MRI and functional MRI; that is, build multimodal based brain templates, because each modality can provide more detailed complementary information from different tissue types (Oishi et al., 2019). Until now, many available infant brain atlases have used the parcellation obtained from the adult brain and propagated onto the infant brain. It is important to obtain more accurate tissue segmentation and parcellation of infant brains by using data from large cohorts of infants and deep learning-based models (Makropoulos et al., 2018; Adamson et al., 2020; Sun et al., 2021). In this study, we have created a set of group-averaged templates, which can be used to determine the average shape and size of the early infant brain. As a result of averaging, however, the sharpness of the image contrast can be lost, and that may be needed when performing highly deformable nonlinear transformations to register images to the template. The template sharpness could be improved or a single-subject template with the size and shape adjusted to the group averaged template could be constructed for building sharp brain templates (Zhao et al., 2019; Oishi et al., 2011). It is noticed that the bilateral thalamic regions surrounding the mid-brain show brighter intensity compared to other subcortical regions from the T1w images in our study. This pattern is consistent with other reported studies (Zhang et al., 2019). It is worth the effort to further investigate the optimization of the acquisition protocol (Zhang et al., 2019).

As normalization-based image analysis is one of the most effective methods for quantification and comparison of the neuroimaging derived measures between populations, an unbiased and representative standard template is critical for the accuracy of the analyses. Future applications of the template will include investigations of the effects of preterm birth and other important perinatal factors, clinical applications such as determining imaging biomarkers for neurological disorders in Chinese populations, and genetic and cultural influences on the brain.

