## Supplemental materials for "Brain Templates for Chinese Babies from Newborn to Three Months of Age"

Table S1. Comparison of average Jacobian determinant of the transformation fields across 90 ROIs when transforming individuals to the unbiased population specific template CUHK_Neo and to population mismatched template UNC_Neo. The majority of the regions show smaller Jacobian determinants when transforming to the population specific template. Only Cingulum_Ant_L, Heschl_L and Heschl_R present significant larger Jacobian determinant when using CUHK_Neo template.

| ROIs | CUHK_NEO | | UNC_NEO | | p values | * if corrected p<0.05 |
|---|---|---|---|---|---|---|
| | mean | stdev | mean | stdev | | |
| Precentral_L | 0.13 | 0.01 | 0.17 | 0.01 | 0.0000 | * |
| Precentral_R | 0.13 | 0.01 | 0.17 | 0.01 | 0.0000 | * |
| Frontal_Sup_L | 0.11 | 0.01 | 0.15 | 0.01 | 0.0000 | * |
| Frontal_Sup_R | 0.11 | 0.00 | 0.15 | 0.01 | 0.0000 | * |
| Frontal_Sup_Orb_L | 0.10 | 0.01 | 0.18 | 0.02 | 0.0000 | * |
| Frontal_Sup_Orb_R | 0.10 | 0.01 | 0.15 | 0.01 | 0.0000 | * |
| Frontal_Mid_L | 0.11 | 0.01 | 0.15 | 0.01 | 0.0000 | * |
| Frontal_Mid_R | 0.11 | 0.00 | 0.15 | 0.01 | 0.0000 | * |
| Frontal_Mid_Orb_L | 0.10 | 0.01 | 0.19 | 0.02 | 0.0000 | * |
| Frontal_Mid_Orb_R | 0.10 | 0.01 | 0.18 | 0.02 | 0.0000 | * |
| Frontal_Inf_Oper_L | 0.15 | 0.01 | 0.17 | 0.02 | 0.0005 | * |
| Frontal_Inf_Oper_R | 0.15 | 0.01 | 0.17 | 0.02 | 0.0008 | |
| Frontal_Inf_Tri_L | 0.13 | 0.01 | 0.16 | 0.01 | 0.0000 | * |
| Frontal_Inf_Tri_R | 0.13 | 0.01 | 0.16 | 0.01 | 0.0000 | * |
| Frontal_Inf_Orb_L | 0.13 | 0.01 | 0.19 | 0.01 | 0.0000 | * |
| Frontal_Inf_Orb_R | 0.13 | 0.01 | 0.18 | 0.01 | 0.0000 | * |
| Rolandic_Oper_L | 0.18 | 0.01 | 0.17 | 0.02 | 0.2026 | |
| Rolandic_Oper_R | 0.16 | 0.01 | 0.16 | 0.02 | 0.9401 | |
| Supp_Motor_Area_L | 0.17 | 0.01 | 0.17 | 0.01 | 0.7320 | |
| Supp_Motor_Area_R | 0.17 | 0.01 | 0.18 | 0.01 | 0.0844 | |
| Olfactory_L | 0.17 | 0.01 | 0.18 | 0.01 | 0.0561 | |
| Olfactory_R | 0.17 | 0.01 | 0.19 | 0.01 | 0.0128 | |
| Frontal_Sup_Medial_L | 0.15 | 0.01 | 0.15 | 0.01 | 0.1790 | |
| Frontal_Sup_Medial_R | 0.13 | 0.01 | 0.16 | 0.01 | 0.0000 | * |
| Frontal_Med_Orb_L | 0.14 | 0.01 | 0.16 | 0.01 | 0.0152 | |
| Frontal_Med_Orb_R | 0.16 | 0.01 | 0.15 | 0.01 | 0.0576 | |
| Rectus_L | 0.13 | 0.01 | 0.17 | 0.01 | 0.0000 | * |
| Rectus_R | 0.13 | 0.01 | 0.16 | 0.01 | 0.0000 | * |
| Insula_L | 0.16 | 0.01 | 0.17 | 0.01 | 0.2064 | |
| Insula_R | 0.16 | 0.01 | 0.17 | 0.01 | 0.0418 | |
| Cingulum_Ant_L | 0.22 | 0.01 | 0.19 | 0.01 | 0.0001 | * |

| | | | | | | |
|---|---|---|---|---|---|---|
| Cingulum_Ant_R | 0.21 | 0.02 | 0.21 | 0.01 | 0.4929 | |
| Cingulum_Mid_L | 0.20 | 0.01 | 0.19 | 0.01 | 0.0024 | |
| Cingulum_Mid_R | 0.20 | 0.01 | 0.20 | 0.01 | 0.8192 | |
| Cingulum_Post_L | 0.20 | 0.01 | 0.22 | 0.02 | 0.0173 | |
| Cingulum_Post_R | 0.20 | 0.02 | 0.19 | 0.02 | 0.5005 | |
| Hippocampus_L | 0.17 | 0.01 | 0.22 | 0.02 | 0.0000 | * |
| Hippocampus_R | 0.16 | 0.01 | 0.21 | 0.01 | 0.0000 | * |
| ParaHippocampal_L | 0.15 | 0.01 | 0.22 | 0.02 | 0.0000 | * |
| ParaHippocampal_R | 0.15 | 0.01 | 0.22 | 0.03 | 0.0000 | * |
| Amygdala_L | 0.14 | 0.01 | 0.22 | 0.02 | 0.0000 | * |
| Amygdala_R | 0.14 | 0.01 | 0.24 | 0.02 | 0.0000 | * |
| Calcarine_L | 0.19 | 0.02 | 0.20 | 0.02 | 0.0950 | |
| Calcarine_R | 0.20 | 0.02 | 0.19 | 0.02 | 0.8549 | |
| Cuneus_L | 0.15 | 0.01 | 0.15 | 0.01 | 0.6279 | |
| Cuneus_R | 0.15 | 0.01 | 0.15 | 0.01 | 0.7536 | |
| Lingual_L | 0.17 | 0.01 | 0.21 | 0.01 | 0.0000 | * |
| Lingual_R | 0.17 | 0.00 | 0.22 | 0.01 | 0.0000 | * |
| Occipital_Sup_L | 0.12 | 0.01 | 0.13 | 0.01 | 0.0024 | |
| Occipital_Sup_R | 0.13 | 0.01 | 0.14 | 0.01 | 0.0351 | |
| Occipital_Mid_L | 0.12 | 0.01 | 0.13 | 0.01 | 0.0005 | * |
| Occipital_Mid_R | 0.11 | 0.01 | 0.12 | 0.02 | 0.0143 | |
| Occipital_Inf_L | 0.10 | 0.02 | 0.16 | 0.02 | 0.0000 | * |
| Occipital_Inf_R | 0.11 | 0.01 | 0.17 | 0.03 | 0.0000 | * |
| Fusiform_L | 0.13 | 0.01 | 0.16 | 0.01 | 0.0000 | * |
| Fusiform_R | 0.13 | 0.01 | 0.16 | 0.01 | 0.0000 | * |
| Postcentral_L | 0.15 | 0.01 | 0.16 | 0.01 | 0.0143 | |
| Postcentral_R | 0.16 | 0.01 | 0.16 | 0.01 | 0.4784 | |
| Parietal_Sup_L | 0.11 | 0.01 | 0.13 | 0.01 | 0.0000 | * |
| Parietal_Sup_R | 0.10 | 0.01 | 0.12 | 0.01 | 0.0000 | * |
| Parietal_Inf_L | 0.12 | 0.01 | 0.13 | 0.01 | 0.0028 | |
| Parietal_Inf_R | 0.12 | 0.00 | 0.12 | 0.01 | 0.3472 | |
| SupraMarginal_L | 0.12 | 0.01 | 0.16 | 0.01 | 0.0000 | * |
| SupraMarginal_R | 0.12 | 0.01 | 0.15 | 0.01 | 0.0000 | * |
| Angular_L | 0.11 | 0.01 | 0.13 | 0.01 | 0.0004 | * |
| Angular_R | 0.11 | 0.01 | 0.12 | 0.01 | 0.0048 | |
| Precuneus_L | 0.16 | 0.01 | 0.16 | 0.01 | 0.0971 | |
| Precuneus_R | 0.17 | 0.01 | 0.17 | 0.01 | 0.7722 | |
| Paracentral_Lobule_L | 0.18 | 0.01 | 0.17 | 0.01 | 0.3990 | |
| Paracentral_Lobule_R | 0.16 | 0.02 | 0.18 | 0.01 | 0.0065 | |
| Caudate_L | 0.20 | 0.01 | 0.22 | 0.02 | 0.0182 | |
| Caudate_R | 0.19 | 0.01 | 0.23 | 0.01 | 0.0000 | * |
| Putamen_L | 0.17 | 0.01 | 0.18 | 0.01 | 0.0032 | |
| Putamen_R | 0.17 | 0.02 | 0.20 | 0.01 | 0.0002 | * |
| Pallidum_L | 0.17 | 0.02 | 0.27 | 0.03 | 0.0000 | * |
| Pallidum_R | 0.17 | 0.02 | 0.32 | 0.04 | 0.0000 | * |
| Thalamus_L | 0.17 | 0.01 | 0.18 | 0.02 | 0.2034 | |
| Thalamus_R | 0.17 | 0.02 | 0.19 | 0.02 | 0.0090 | |
| Heschl_L | 0.23 | 0.03 | 0.17 | 0.02 | 0.0001 | * |
| Heschl_R | 0.28 | 0.03 | 0.18 | 0.04 | 0.0000 | * |
| Temporal_Sup_L | 0.16 | 0.01 | 0.17 | 0.01 | 0.0127 | |
| Temporal_Sup_R | 0.15 | 0.01 | 0.16 | 0.01 | 0.0003 | * |

| | | | | | | |
|---|---|---|---|---|---|---|
| Temporal_Pole_Sup_L | 0.18 | 0.01 | 0.18 | 0.01 | 0.5703 | |
| Temporal_Pole_Sup_R | 0.17 | 0.01 | 0.18 | 0.01 | 0.0963 | |
| Temporal_Mid_L | 0.11 | 0.01 | 0.15 | 0.01 | 0.0000 | * |
| Temporal_Mid_R | 0.12 | 0.00 | 0.14 | 0.01 | 0.0000 | * |
| Temporal_Pole_Mid_L | 0.12 | 0.02 | 0.18 | 0.02 | 0.0000 | * |
| Temporal_Pole_Mid_R | 0.11 | 0.01 | 0.18 | 0.01 | 0.0000 | * |
| Temporal_Inf_L | 0.10 | 0.01 | 0.14 | 0.00 | 0.0000 | * |
| Temporal_Inf_R | 0.10 | 0.00 | 0.14 | 0.00 | 0.0000 | * |

Table S2. Comparison of the standard deviation of the registered images across 90 ROIs when transforming individuals to the unbiased population specific template CUHK_Neo and to population mismatched template UNC_Neo. The majority of the regions show smaller standard deviation (16 of them were significant after Bonferroni correction) when using the population specific template.

| **ROIs** | **CUHK_NEO** | | **UNC_NEO** | | **p values** | *** if corrected p<0.05** |
|---|---|---|---|---|---|---|
| | **mean** | **stdev** | **mean** | **stdev** | | |
| Precentral_L | 5.76 | 0.15 | 6.03 | 0.12 | 0.0003 | * |
| Precentral_R | 5.89 | 0.22 | 6.26 | 0.24 | 0.0021 | |
| Frontal_Sup_L | 6.19 | 0.19 | 6.51 | 0.18 | 0.0010 | |
| Frontal_Sup_R | 6.17 | 0.28 | 6.54 | 0.23 | 0.0050 | |
| Frontal_Sup_Orb_L | 4.86 | 0.25 | 5.13 | 0.26 | 0.0322 | |
| Frontal_Sup_Orb_R | 4.99 | 0.21 | 5.30 | 0.26 | 0.0090 | |
| Frontal_Mid_L | 5.66 | 0.21 | 5.96 | 0.19 | 0.0036 | |
| Frontal_Mid_R | 5.57 | 0.25 | 5.93 | 0.22 | 0.0027 | |
| Frontal_Mid_Orb_L | 5.91 | 0.21 | 6.29 | 0.30 | 0.0039 | |
| Frontal_Mid_Orb_R | 5.81 | 0.32 | 6.15 | 0.44 | 0.0614 | |
| Frontal_Inf_Oper_L | 5.51 | 0.20 | 5.68 | 0.25 | 0.1008 | |
| Frontal_Inf_Oper_R | 5.42 | 0.36 | 5.64 | 0.37 | 0.1965 | |
| Frontal_Inf_Tri_L | 5.43 | 0.16 | 5.83 | 0.15 | 0.0000 | * |
| Frontal_Inf_Tri_R | 5.39 | 0.31 | 5.78 | 0.25 | 0.0058 | |
| Frontal_Inf_Orb_L | 5.60 | 0.25 | 6.08 | 0.31 | 0.0014 | |
| Frontal_Inf_Orb_R | 5.60 | 0.23 | 6.09 | 0.25 | 0.0003 | * |
| Rolandic_Oper_L | 5.94 | 0.13 | 5.99 | 0.19 | 0.5107 | |
| Rolandic_Oper_R | 5.51 | 0.25 | 5.67 | 0.28 | 0.2048 | |
| Supp_Motor_Area_L | 6.52 | 0.33 | 6.66 | 0.22 | 0.2806 | |
| Supp_Motor_Area_R | 6.32 | 0.36 | 6.45 | 0.25 | 0.3350 | |
| Olfactory_L | 5.60 | 0.25 | 5.97 | 0.31 | 0.0085 | |
| Olfactory_R | 6.19 | 0.26 | 7.36 | 0.48 | 0.0000 | * |
| Frontal_Sup_Medial_L | 5.84 | 0.20 | 6.14 | 0.20 | 0.0043 | |
| Frontal_Sup_Medial_R | 6.16 | 0.32 | 6.66 | 0.21 | 0.0005 | * |
| Frontal_Med_Orb_L | 5.54 | 0.19 | 5.91 | 0.33 | 0.0060 | |
| Frontal_Med_Orb_R | 5.29 | 0.18 | 5.75 | 0.28 | 0.0003 | * |
| Rectus_L | 5.46 | 0.36 | 6.16 | 0.45 | 0.0013 | |
| Rectus_R | 5.30 | 0.33 | 5.87 | 0.29 | 0.0008 | |
| Insula_L | 4.16 | 0.24 | 4.24 | 0.23 | 0.4503 | |
| Insula_R | 3.94 | 0.27 | 4.09 | 0.27 | 0.2294 | |
| Cingulum_Ant_L | 5.33 | 0.13 | 5.79 | 0.16 | 0.0000 | * |
| Cingulum_Ant_R | 4.83 | 0.22 | 5.19 | 0.21 | 0.0018 | |
| Cingulum_Mid_L | 5.65 | 0.53 | 5.72 | 0.51 | 0.7754 | |
| Cingulum_Mid_R | 5.42 | 0.49 | 5.60 | 0.49 | 0.4122 | |
| Cingulum_Post_L | 6.71 | 0.52 | 7.74 | 0.41 | 0.0001 | * |

| | | | | | | |
|---|---|---|---|---|---|---|
| Cingulum_Post_R | 6.72 | 0.70 | 7.33 | 0.75 | 0.0737 | |
| Hippocampus_L | 5.01 | 0.30 | 6.07 | 0.33 | 0.0000 | * |
| Hippocampus_R | 4.81 | 0.37 | 5.54 | 0.37 | 0.0003 | * |
| ParaHippocampal_L | 6.74 | 0.29 | 8.80 | 0.60 | 0.0000 | * |
| ParaHippocampal_R | 6.68 | 0.24 | 8.31 | 0.47 | 0.0000 | * |
| Amygdala_L | 3.84 | 0.25 | 4.91 | 0.36 | 0.0000 | * |
| Amygdala_R | 3.88 | 0.37 | 5.10 | 0.50 | 0.0000 | * |
| Calcarine_L | 5.85 | 0.25 | 6.07 | 0.32 | 0.1037 | |
| Calcarine_R | 5.30 | 0.46 | 5.50 | 0.47 | 0.3485 | |
| Cuneus_L | 5.36 | 0.29 | 5.53 | 0.35 | 0.2445 | |
| Cuneus_R | 5.50 | 0.38 | 5.68 | 0.40 | 0.3213 | |
| Lingual_L | 5.73 | 0.21 | 5.99 | 0.23 | 0.0146 | |
| Lingual_R | 6.12 | 0.27 | 6.68 | 0.36 | 0.0010 | |
| Occipital_Sup_L | 5.23 | 0.34 | 5.23 | 0.37 | 0.9849 | |
| Occipital_Sup_R | 5.04 | 0.34 | 5.12 | 0.29 | 0.5525 | |
| Occipital_Mid_L | 4.68 | 0.32 | 4.81 | 0.34 | 0.3656 | |
| Occipital_Mid_R | 4.94 | 0.29 | 5.06 | 0.31 | 0.3895 | |
| Occipital_Inf_L | 5.33 | 0.59 | 5.40 | 0.59 | 0.7905 | |
| Occipital_Inf_R | 5.32 | 0.37 | 5.41 | 0.42 | 0.6177 | |
| Fusiform_L | 5.62 | 0.12 | 5.95 | 0.13 | 0.0000 | * |
| Fusiform_R | 5.53 | 0.21 | 5.80 | 0.19 | 0.0076 | |
| Postcentral_L | 6.06 | 0.25 | 6.40 | 0.27 | 0.0087 | |
| Postcentral_R | 5.91 | 0.26 | 6.29 | 0.33 | 0.0102 | |
| Parietal_Sup_L | 6.08 | 0.35 | 6.29 | 0.33 | 0.1767 | |
| Parietal_Sup_R | 6.80 | 0.39 | 7.05 | 0.43 | 0.1866 | |
| Parietal_Inf_L | 5.41 | 0.22 | 5.52 | 0.22 | 0.2672 | |
| Parietal_Inf_R | 5.30 | 0.27 | 5.41 | 0.32 | 0.4302 | |
| SupraMarginal_L | 5.77 | 0.33 | 5.98 | 0.29 | 0.1458 | |
| SupraMarginal_R | 5.43 | 0.25 | 5.53 | 0.22 | 0.3250 | |
| Angular_L | 4.79 | 0.25 | 4.90 | 0.28 | 0.3498 | |
| Angular_R | 5.03 | 0.22 | 5.14 | 0.25 | 0.3304 | |
| Precuneus_L | 5.92 | 0.22 | 6.09 | 0.23 | 0.1055 | |
| Precuneus_R | 5.64 | 0.35 | 5.85 | 0.37 | 0.2225 | |
| Paracentral_Lobule_L | 6.83 | 0.33 | 6.98 | 0.33 | 0.3167 | |
| Paracentral_Lobule_R | 6.01 | 0.43 | 6.29 | 0.40 | 0.1518 | |
| Caudate_L | 4.72 | 0.48 | 5.09 | 0.48 | 0.1062 | |
| Caudate_R | 4.12 | 0.58 | 4.43 | 0.60 | 0.2502 | |
| Putamen_L | 3.49 | 0.57 | 3.64 | 0.51 | 0.5309 | |
| Putamen_R | 3.26 | 0.53 | 3.48 | 0.57 | 0.3905 | |
| Pallidum_L | 4.20 | 0.75 | 4.48 | 0.53 | 0.3383 | |
| Pallidum_R | 4.05 | 0.71 | 4.18 | 0.65 | 0.6811 | |
| Thalamus_L | 5.12 | 0.90 | 5.25 | 0.77 | 0.7315 | |
| Thalamus_R | 4.94 | 0.95 | 4.96 | 0.97 | 0.9607 | |
| Heschl_L | 6.00 | 0.27 | 6.12 | 0.29 | 0.3592 | |
| Heschl_R | 6.51 | 0.54 | 6.69 | 0.42 | 0.4170 | |
| Temporal_Sup_L | 6.28 | 0.21 | 6.53 | 0.21 | 0.0202 | |
| Temporal_Sup_R | 6.01 | 0.30 | 6.25 | 0.30 | 0.1002 | |
| Temporal_Pole_Sup_L | 8.07 | 0.41 | 8.83 | 0.38 | 0.0004 | * |
| Temporal_Pole_Sup_R | 8.26 | 0.56 | 8.84 | 0.53 | 0.0305 | |
| Temporal_Mid_L | 5.21 | 0.16 | 5.41 | 0.16 | 0.0122 | |
| Temporal_Mid_R | 5.21 | 0.29 | 5.37 | 0.26 | 0.1931 | |

| | | | | | |
|---|---|---|---|---|---|
| Temporal_Pole_Mid_L | 7.90 | 0.48 | 7.96 | 0.48 | 0.7804 |
| Temporal_Pole_Mid_R | 7.40 | 0.47 | 7.45 | 0.40 | 0.8105 |
| Temporal_Inf_L | 5.56 | 0.23 | 5.78 | 0.23 | 0.0551 |
| Temporal_Inf_R | 5.65 | 0.24 | 5.87 | 0.22 | 0.0438 |

Table S3. Age and sex effect on regional brain volumes. With the GLM: volume ~ Gestational Age + Corrected Age (days) + Sex, the uncorrected and Bonferroni corrected ps for corrected age at scan and sex were reported for 90 ROIs.

| **ROIs** | **p for Sex** | | **p for Age** | |
|---|---|---|---|---|
| | **P values** | *** if corrected p<0.05** | **P values** | *** if corrected p<0.05** |
| Precentral_L | 0.0007 | | 0.0000 | * |
| Precentral_R | 0.0001 | * | 0.0000 | * |
| Frontal_Sup_L | 0.0000 | * | 0.0000 | * |
| Frontal_Sup_R | 0.0018 | | 0.0000 | * |
| Frontal_Sup_Orb_L | 0.0006 | | 0.0000 | * |
| Frontal_Sup_Orb_R | 0.0000 | * | 0.0000 | * |
| Frontal_Mid_L | 0.0011 | | 0.0000 | * |
| Frontal_Mid_R | 0.0006 | | 0.0000 | * |
| Frontal_Mid_Orb_L | 0.0006 | | 0.0000 | * |
| Frontal_Mid_Orb_R | 0.0042 | | 0.0000 | * |
| Frontal_Inf_Oper_L | 0.0072 | | 0.0000 | * |
| Frontal_Inf_Oper_R | 0.0123 | | 0.0000 | * |
| Frontal_Inf_Tri_L | 0.0015 | | 0.0000 | * |
| Frontal_Inf_Tri_R | 0.0015 | | 0.0000 | * |
| Frontal_Inf_Orb_L | 0.0021 | | 0.0000 | * |
| Frontal_Inf_Orb_R | 0.0041 | | 0.0000 | * |
| Rolandic_Oper_L | 0.0033 | | 0.0000 | * |
| Rolandic_Oper_R | 0.0107 | | 0.0000 | * |
| Supp_Motor_Area_L | 0.0006 | | 0.0000 | * |
| Supp_Motor_Area_R | 0.0116 | | 0.0000 | * |
| Olfactory_L | 0.0064 | | 0.0000 | * |
| Olfactory_R | 0.0370 | | 0.0000 | * |
| Frontal_Sup_Medial_L | 0.0021 | | 0.0000 | * |
| Frontal_Sup_Medial_R | 0.0269 | | 0.0000 | * |
| Frontal_Med_Orb_L | 0.0017 | | 0.0000 | * |
| Frontal_Med_Orb_R | 0.0003 | * | 0.0000 | * |
| Rectus_L | 0.0001 | * | 0.0000 | * |
| Rectus_R | 0.0000 | * | 0.0000 | * |
| Insula_L | 0.0004 | * | 0.0000 | * |
| Insula_R | 0.0018 | | 0.0000 | * |
| Cingulum_Ant_L | 0.0148 | | 0.0000 | * |
| Cingulum_Ant_R | 0.0079 | | 0.0000 | * |
| Cingulum_Mid_L | 0.0001 | * | 0.0000 | * |
| Cingulum_Mid_R | 0.0007 | | 0.0000 | * |
| Cingulum_Post_L | 0.0020 | | 0.7338 | |
| Cingulum_Post_R | 0.0576 | | 0.2501 | |
| Hippocampus_L | 0.0284 | | 0.0000 | * |
| Hippocampus_R | 0.0107 | | 0.0000 | * |
| ParaHippocampal_L | 0.0051 | | 0.0000 | * |
| ParaHippocampal_R | 0.0090 | | 0.0000 | * |

| | | | | |
|---|---|---|---|---|
| Amygdala_L | 0.0504 | | 0.0000 | * |
| Amygdala_R | 0.0233 | | 0.0000 | * |
| Calcarine_L | 0.0654 | | 0.0000 | * |
| Calcarine_R | 0.0850 | | 0.0000 | * |
| Cuneus_L | 0.0217 | | 0.0000 | * |
| Cuneus_R | 0.0177 | | 0.0000 | * |
| Lingual_L | 0.0040 | | 0.0000 | * |
| Lingual_R | 0.0084 | | 0.0000 | * |
| Occipital_Sup_L | 0.0453 | | 0.0000 | * |
| Occipital_Sup_R | 0.0024 | | 0.0000 | * |
| Occipital_Mid_L | 0.0069 | | 0.0000 | * |
| Occipital_Mid_R | 0.0014 | | 0.0000 | * |
| Occipital_Inf_L | 0.0018 | | 0.0001 | * |
| Occipital_Inf_R | 0.0646 | | 0.0000 | * |
| Fusiform_L | 0.0023 | | 0.0000 | * |
| Fusiform_R | 0.0241 | | 0.0000 | * |
| Postcentral_L | 0.0238 | | 0.0000 | * |
| Postcentral_R | 0.0020 | | 0.0000 | * |
| Parietal_Sup_L | 0.0333 | | 0.0000 | * |
| Parietal_Sup_R | 0.0251 | | 0.0000 | * |
| Parietal_Inf_L | 0.0048 | | 0.0000 | * |
| Parietal_Inf_R | 0.0709 | | 0.0000 | * |
| SupraMarginal_L | 0.0127 | | 0.0000 | * |
| SupraMarginal_R | 0.0018 | | 0.0000 | * |
| Angular_L | 0.0214 | | 0.0000 | * |
| Angular_R | 0.0013 | | 0.0000 | * |
| Precuneus_L | 0.0056 | | 0.0000 | * |
| Precuneus_R | 0.0043 | | 0.0000 | * |
| Paracentral_Lobule_L | 0.0002 | * | 0.0000 | * |
| Paracentral_Lobule_R | 0.3289 | | 0.0000 | * |
| Caudate_L | 0.0787 | | 0.2160 | |
| Caudate_R | 0.1488 | | 0.0476 | |
| Putamen_L | 0.0454 | | 0.0000 | * |
| Putamen_R | 0.1084 | | 0.0000 | * |
| Pallidum_L | 0.6905 | | 0.0000 | * |
| Pallidum_R | 0.4809 | | 0.0000 | * |
| Thalamus_L | 0.6825 | | 0.0000 | * |
| Thalamus_R | 0.2706 | | 0.0000 | * |
| Heschl_L | 0.0156 | | 0.0000 | * |
| Heschl_R | 0.0335 | | 0.0000 | * |
| Temporal_Sup_L | 0.0401 | | 0.0000 | * |
| Temporal_Sup_R | 0.0089 | | 0.0000 | * |
| Temporal_Pole_Sup_L | 0.0235 | | 0.0000 | * |
| Temporal_Pole_Sup_R | 0.1985 | | 0.0000 | * |
| Temporal_Mid_L | 0.0404 | | 0.0000 | * |
| Temporal_Mid_R | 0.0064 | | 0.0000 | * |
| Temporal_Pole_Mid_L | 0.0050 | | 0.0000 | * |
| Temporal_Pole_Mid_R | 0.0823 | | 0.0000 | * |
| Temporal_Inf_L | 0.0034 | | 0.0000 | * |
| Temporal_Inf_R | 0.0023 | | 0.0000 | * |
